# Phospholipid Metabolism Is Associated with HIV Rebound Upon Treatment Interruption

**DOI:** 10.1101/2020.12.09.418814

**Authors:** Leila B. Giron, Emmanouil Papasavvas, Xiangfan Yin, Aaron R. Goldman, Hsin-Yao Tang, Clovis S. Palmer, Alan L. Landay, Jonathan Z. Li, John R. Koethe, Karam Mounzer, Jay R. Kostman, Qin Liu, Luis J. Montaner, Mohamed Abdel-Mohsen

## Abstract

Lipids are biologically active molecules involved in a variety of cellular processes and immunological functions, including inflammation. It was recently shown that phospholipids and their derivatives, lysophospholipids, can reactivate latent (dormant) tumor cells, causing cancer recurrence. However, the potential link between lipids and HIV latency, persistence, and viral rebound after cessation of antiretroviral therapy (ART) has never been investigated. We explored the links between plasma lipids and the burden of HIV during ART. We profiled the circulating lipidome from the plasma of 24 chronically HIV-infected individuals on suppressive ART who subsequently underwent an analytic treatment interruption (ATI) without concurrent immunotherapies. The pre-ATI viral burden was estimated as time-to-viral-rebound and viral load setpoints post-ATI. We found that higher pre-ATI levels of lysophospholipids, including the pro-inflammatory lysophosphatidylcholine, were associated with faster time-to-viral-rebound and higher viral setpoints upon ART cessation. Furthermore, higher pre-ATI levels of the pro-inflammatory byproduct of intestinal lysophosphatidylcholine metabolism, trimethylamine-N-oxide (TMAO), were also linked to faster viral rebound post-ART. Finally, pre-ATI levels of several phosphatidylcholine species (lysophosphatidylcholine precursors) correlated strongly with higher pre-ATI levels of HIV DNA in peripheral CD4+ T cells. Our proof-of-concept data point to phospholipids and lysophospholipids as plausible pro-inflammatory contributors to HIV persistence and rapid post-ART HIV rebound. The potential interplay between phospholipid metabolism and both the establishment and maintenance of HIV latent reservoirs during- and post-ART warrants further investigation.

**IMPORTANCE:** The likelihood of HIV rebound after stopping antiretroviral therapy (ART) is a function of the interplay between the size of HIV reservoirs that persist despite ART and the host immunological and inflammatory factors that modulate these reservoirs. There is a need to comprehensively understand these host factors to develop a strategy to cure HIV infection and prevent viral rebound post-ART. Lipids are important biologically active molecules that are known to mediate several cellular functions, including reactivating latent tumor cells; however, their role in HIV latency, persistence, and post-ART rebound has never been investigated. We observed significant links between higher levels of the pro-inflammatory lysophosphatidylcholine and its intestinal metabolic byproduct, trimethylamine-N-oxide, and both faster time-to-viral rebound and higher viral load setpoint post ART. These data highlight the need for further studies to understand the potential contribution of phosphatidylcholine and lysophosphatidylcholine metabolism in shaping host immunological and inflammatory milieu during- and post-ART.

## TEXT

A comprehensive understanding of the host factors modulating HIV persistence is imperative for developing effective strategies to eradicate the latent HIV reservoir, which persists despite antiretroviral therapy (ART) and causes viral rebound upon ART discontinuation (1). Lipids are biologically active molecules involved in a broad range of cellular processes and immunological functions, including inflammation (2, 3). It was recently shown that phospholipids and their derivatives, lysophospholipids, can reactivate latent (dormant) tumor cells, causing cancer recurrence (4). While the interplay between lipids and both HIV and ART has been studied in the context of the development of inflammation-associated comorbidities, particularly subclinical atherosclerosis (5-9), the potential impact of lipids on HIV latency, persistence, and post-ART rebound has never been investigated.

There is currently no standard method to measure the total body burden of the replication-competent HIV reservoir (1, 10). However, a possible way to estimate both the overall size of the HIV reservoir and the degree of viral control is by assessing time-to-viral-rebound and/or viral load setpoints upon cessation of ART. In this study, we profiled the circulating lipidome from the plasma of 24 chronically HIV-infected individuals on suppressive ART who subsequently underwent an analytic treatment interruption (ATI) (11, 12). All 24 individuals underwent ATI without concurrent immunomodulatory agents that might confound our analysis. Lipidomic analysis was performed using liquid chromatography–mass spectrometry (LC–MS), as described previously (13), on plasma samples collected immediately before ATI. Both time-to-viral-rebound and viral load setpoints were measured during ATI. This cohort had a wide distribution of viral rebound times (14 to 119 days; median=28) and viral load setpoints (median=13,675 copies/ml; **Supplementary Table 1**). Using these data, we investigated whether there is a link between pre-ATI lipid profiles and the body burden of HIV during ART (estimated as post-ATI time-to-viral-rebound and viral load setpoints).

### Levels of plasma lysophospholipids measured pre-ATI associate with time-to-viral-rebound post-ATI

We identified a total of 967 lipids, belong to 21 lipids classes (described in **Supplementary Table 2**), in the plasma samples. Using the Cox proportional-hazards model, we found that pre-ATI levels of several of these lipids significantly associated with a faster time-to-viral-rebound (**Figure 1A;** lipids with a hazard ratio (HR)>5 and *P*<0.01 are labeled). We next examined whether these lipids belong to particular lipidomic pathways or classes. Pathway analysis of all lipids whose pre-ATI levels associated with time-to-viral-rebound with *P*<0.05 showed that the pathway most associated with viral rebound was glycerophospholipid metabolism (**Figure 1B**). The pre-ART levels of three lipid classes were significantly (FDR<0.05) associated with faster time-to-viral-rebound (**Figure 1C** and **Supplementary Table 3**): lysophosphatidylcholine (LPC), lysophosphatidylethanolamine (LPE), and lysophospholipid acid (LPA). All three classes belong to the lysophospholipid group, which is a subgroup of the glycerophospholipid family shown in Figure 1B. Lysophospholipids are small bioactive lipid molecules known to play important roles in regulating several biological functions, including promoting inflammation (8, 14-19). The significant associations between these lysophospholipid classes and faster time-to-viral-rebound were confirmed using two additional, independent analyses: Mantel-Cox survival test, after separating participants into low or high groups based on the median level of each of these lipid classes (**Figure 1D)**; and Spearman’s rank correlation between the levels of these lipid classes and time-to-viral-rebound (**Supplementary Table 3)**. These data point, for the first time, to plausible links between phospholipid and lysophospholipid metabolism and HIV rebound post ART. Intriguingly, similar phospholipids and lysophospholipids were recently shown to reactivate latent (dormant) cancer cells (4). Our exploratory findings, that are consistent with the reported functions of these lysophospholipids, raise the question of whether these lysophospholipids condition the host environment with higher levels of inflammation that might impact viral reactivation, cellular processes, and immunological functions during- and/or post-ATI.

**Figure 1.**
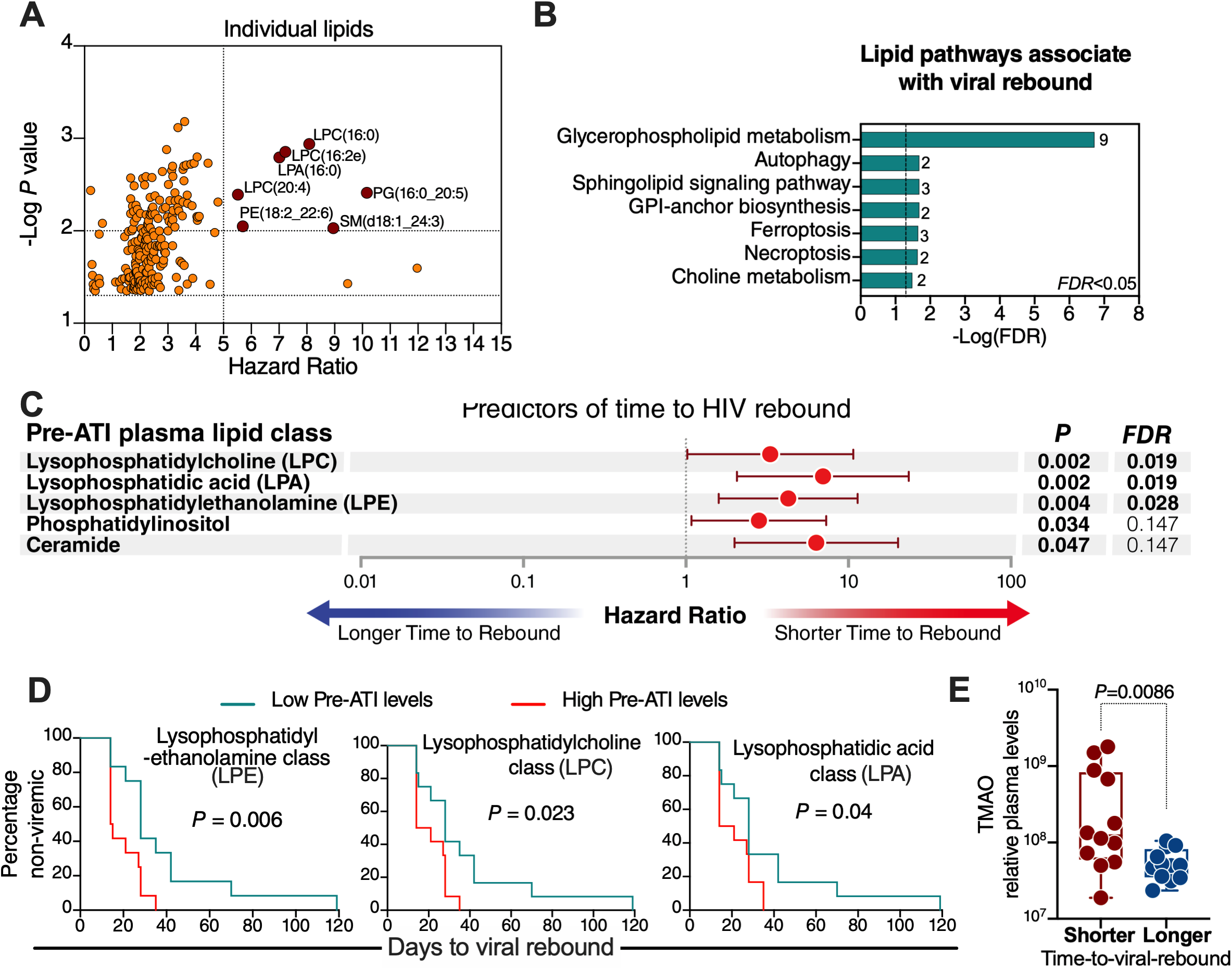
Higher pre-ATI lysophospholipid metabolism and its bioactive byproducts associate with faster post-ATI time-to-viral-rebound. **(A)** Lipids whose pre-ATI levels associate with post-ATI time-to-viral-rebound, as determined by the Cox proportional-hazards model. Lipids with *P*<0.01 and hazard ratio (HR) > 5 are shown in red and labeled. **(B)** Lipid pathway analysis of plasma lipids whose pre-ATI levels associated with time-to-viral-rebound with *P*<0.05 using LIPEA (Lipid Pathway Enrichment Analysis; https://lipea.biotec.tu-dresden.de/home). The graph shows all implicated pathways with FDR<0.05. Numbers beside each pathway represent the number of dysregulated lipids within the particular pathway. **(C)** Lipid classes whose pre-ATI levels associate with post-ATI time-to-viral-rebound, as determined by the Cox proportional-hazards model. FDR was calculated using the Benjamini–Hochberg approach. **(D)** Confirmatory analysis of the three lysophospholipid classes using the Mantel-Cox test. Low pre-ATI levels = lower than group median; High pre-ATI levels = higher than group median. **(E)** Participants were separated into shorter or longer time-to-viral-rebound groups by the median of time-to-viral-rebound; levels of TMAO were higher in individuals who rebounded faster compared to individuals who rebounded slower. Mann–Whitney U test was used for statistical analysis. Statistical analyses were performed in R and Prism 7.0 (GraphPad).

### Levels of plasma trimethylamine-N-oxide (TMAO) measured pre-ATI associate with post-ATI time-to-viral-rebound

The pro-inflammatory lipid class LPC can be hydrolyzed in the intestine to LPA and choline; choline can be metabolized into trimethylamine, which is converted to trimethylamine-N-oxide (TMAO) in the liver (20). TMAO induces several pro-inflammatory mediators and has been implicated in several inflammation-associated diseases (20-23). Given that LPC and LPA lipids were among the lipids whose pre-ATI levels associated with faster viral rebound upon ART cessation (**Figure 1A-D**), we sought to examine if levels of TMAO associated with post-ATI time-to-viral rebound. We performed metabolomics analysis, using LC-MS, as described previously (24), on the same pre-ATI plasma samples. Indeed, pre-ATI levels of TMAO were higher in individuals with lower than the median days to viral rebound (fast rebounders) compared to individuals with higher than the median days to rebound (delayed rebounders) (**Figure 1E**). Our observations that the pro-inflammatory byproducts of intestinal LPC metabolism (LPA and TMAO) are also associated with faster HIV rebound demand a greater understanding of the interaction between glycerophospholipid or choline metabolism by intestinal microbiota and viral persistence during-ART or rebound post-ART. Such understanding may inform therapeutic approaches targeting the gut microbiota–lipid metabolism interface to reduce inflammation and facilitate the clearance of HIV reservoirs.

### Pre-ATI plasma lysophospholipids associate with post-ATI viral load setpoint

In addition to time-to-viral-rebound, post-ART viral load setpoint can reflect the body burden of HIV during ART. Therefore, we asked whether pre-ATI lipid profiles are associated with post-ATI viral load setpoint. Pre-ATI levels of several lipids associated with post-ATI viral load setpoint with *P<*0.01 and Spearman *rho*>0.5 (**Figure 2A**). Furthermore, pre-ATI LPC and LPE classes levels correlated with post-ATI viral load setpoint (**Figure 2B-C**, respectively). Finally, the pre-ATI levels of the LPC (24:0) lipid species, which was one of the individual lipids whose pre-ATI level correlated with time-to-viral-rebound (**Figure 1A**), also associated with post-ATI viral load setpoints (**Figure 2D**). Notably, levels of LPC (20:4) during HIV infection have been shown to associate with the progression of carotid artery atherosclerosis, even after ART suppression (6). These data indicate that pre-ATI phospholipid metabolism is linked to viral load setpoint upon ART cessation.

**Figure 2.**
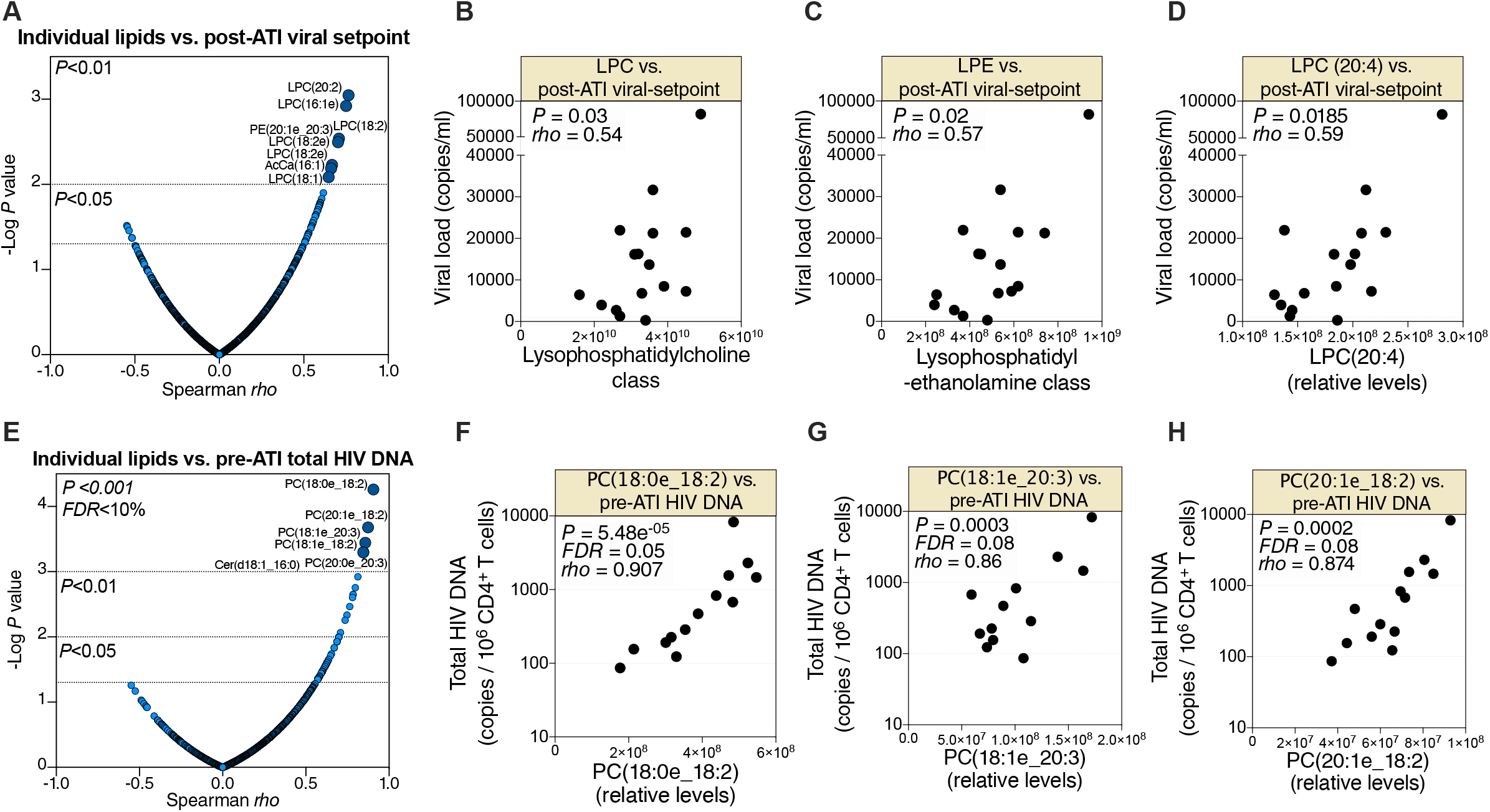
Pre-ATI phospholipid metabolism associates with post-ATI viral load setpoint and pre-ATI HIV DNA. **(A)** Spearman’s rank correlations between pre-ATI lipids and post-ATI viral load setpoint. Lipids with *P*<0.01 and Spearman *rho*>0.5 are shown in dark blue and are labeled. **(B-D)** Correlations between pre-ATI levels of LPC class **(B)**, LPE class **(C)**, or LPC (20:4) lipid species **(D)** and post-ATI viral load setpoint. Each dot represents an HIV^+^ individual. **(D)** Spearman’s rank correlations between pre-ATI lipids and pre-ATI total HIV DNA measured in peripheral CD4^+^ T cells. Lipids with *FDR*<0.1 and Spearman *rho*>0.5 are shown in dark blue and are labeled. **(F-H)** Correlations between pre-ATI levels of several phosphatidylcholine species and pre-ATI levels of HIV DNA in peripheral CD4^+^ T cells. All correlations were evaluated using Spearman’s rank correlation coefficient tests. Statistical analyses were performed in R and Prism 7.0 (GraphPad).

### Pre-ATI phosphatidylcholines associate with pre-ATI HIV DNA in the periphery

Finally, we examined the links between pre-ATI plasma lipidome and pre-ATI total HIV DNA measured in periphery CD4^+^ T by droplet digital PCR (ddPCR), as described (25). Levels of several phosphatidylcholine species (precursors of lysophosphatidylcholine) significantly correlated (FDR <10%) with CD4^+^ T cell-associated HIV DNA (**Figure 2E-H**). These data further highlight the potential links between phospholipid metabolism and HIV persistence.

Our exploratory study has limitations, including small sample size and sampling of blood. The sample size did not allow for addressing the confounding effects of age, gender, ethnicity, weight, diet, duration-of-infection, duration-on-ART, ART-regimen, or comorbidities on lipidomic signatures. Addressing the impact of these confounders and validating our data using larger cohorts should be the subject of future studies. In addition, analyzing lipids and HIV burden in different tissues, including adipose tissue, and mechanistic studies *in vitro* and in animal models of HIV infection will be needed to examine the precise interplay between phospholipid metabolism and viral persistence. Such studies might identify lipid-based interactions that can be targeted to decrease the size of HIV reservoirs and/or delay viral rebound after stopping ART.

Despite these limitations, our study provides the first proof-of-concept evidence that phospholipid metabolism might be involved in a host milieu that facilitates a faster HIV rebound after ART cessation. The potential interactions between phospholipid/lysophospholipid metabolism and both the establishment and maintenance of HIV latency warrant further investigation.

## SUPPLEMENTARY MATERIALS

**Supplementary Table 1**. Clinical and demographic data of the study cohort.

**Supplementary Table 2**. 967 lipids identified in this study were assigned to 21 lipid classes.

**Supplementary Table 3**. A list of lipid classes whose pre-ATI levels associate with faster time-to-viral-rebound upon ART cessation.

## AUTHOR CONTRIBUTIONS

M.A-M conceived and designed the study. L.B.G carried out experiments. C.S.P, A.L.L, J.Z.L, J.R.K analyzed and interpreted lipidomic and metabolic data. K.M, J.R.K, E.P, and L.J.M, selected study participants and interpreted clinical data. A.R.G and H.T performed lipidomic and metabolic analysis. X.Y and Q.L performed statistical analysis. L.B.G and M.A-M wrote the manuscript, and all authors edited it.

## ACKNOWLEDGMENTS

This work is supported by the Foundation for AIDS Research (amfAR) impact grant # 109840-65-RGRL to M.A-M and the NIH R21 AI143385 to M.A-M. M.A-M is also supported by NIH grants (R01 DK123733, R01 AG062383, R01NS117458, R21 AI129636, and R21 NS106970), the Penn Center for AIDS Research (P30 AI 045008), and W.W. Smith Charitable Trust grant # A1901. Lipidomic and Metabolomic analyses were performed by the Wistar Proteomics and Metabolomics Shared Resource supported in part by NIH Cancer Center Support Grant CA010815 on a Thermo Q-Exactive HF-X mass spectrometer purchased with NIH grant S10 OD023586. We would like to thank Rachel E. Locke, Ph.D., for providing comments. We would like to thank all donor participants.

